# Breaking down of healthcare system: Mathematical modelling for controlling the novel coronavirus (2019-nCoV) outbreak in Wuhan, China

**DOI:** 10.1101/2020.01.27.922443

**Authors:** Wai-Kit Ming, Jian Huang, Casper J. P. Zhang

## Abstract

**Background:** A novel coronavirus pneumonia initially identified in Wuhan, China and provisionally named 2019-nCoV has surged in the public. In anticipation of substantial burdens on healthcare system following this human-to-human spread, we aim to scrutinise the currently available information and evaluate the burden of healthcare systems during this outbreak in Wuhan.

**Methods and Findings:** We applied a modified SIR model to project the actual number of infected cases and the specific burdens on isolation wards and intensive care units (ICU), given the scenarios of different diagnosis rates as well as different public health intervention efficacy. Our estimates suggest, assuming 50% diagnosis rate if no public health interventions were implemented, that the actual number of infected cases could be much higher than the reported, with estimated 88,075 cases (as of 31^st^ January, 2020), and projected burdens on isolation wards and ICU would be 34,786 and 9,346 respectively The estimated burdens on healthcare system could be largely reduced if at least 70% efficacy of public health intervention is achieved.

**Conclusion:** The health system burdens arising from the actual number of cases infected by the novel coronavirus appear to be considerable if no effective public health interventions were implemented. This calls for continuation of implemented anti-transmission measures (e.g., closure of schools and facilities, suspension of public transport, lockdown of city) and further effective large-scale interventions spanning all subgroups of populations (e.g., universal facemask wear) aiming at obtaining overall efficacy with at least 70% to ensure the functioning of and to avoid the breakdown of health system.

## 1. Background

A novel coronavirus pneumonia, initially identified in Wuhan, Central China and now named as 2019-nCoV[1], has surged in the public. As from late January 2020, authorities had reported more than 6,000 confirmed cases across nearly all provinces in mainland China and confirmed over 130 deaths. Globally, more than 13 countries or regions have reported confirmed cases including domestic cases. With the increasing incidence of confirmed cases, corresponding spread control policies and emergency actions are taking place.

The symptom onset date of the first 2019-nCoV patient was identified in early December 2019 and the outbreak started in late December with most of cases epidemiologically connected to a seafood market in the city of Wuhan, Hubei province, China [2]. Following the cases reported in other Chinese cities and overseas, the National Health Commission (NHC) of People’s Republic of China confirmed the evidence of human-to-human transmission of such viral pneumonia[3]. Most of confirmed cases so far are travellers from or ever been to Wuhan or other Chinese cities. Several counties also reported their first domestic cases. The number of confirmed cases is expected to increase given the availability of fast-track laboratory test and anticipated country-wide commute arising from Chinese new year holidays.

To combat the 2019-nCoV outbreak, authorities in China have implemented several preventive measures. Starting from 10am, 23^rd^ January, all public transport has been temporarily suspended following by the lockdown on the city of Wuhan[4]. Neighbouring cities also announced a lockdown in sequence. Local residents were advised to remain at home and avoid gathering in order to contain the virus spread. Following the raise of protection standards instructed by NHC, prevention and control measures, such as disinfection for public facilities, have been strengthened and taking places in other cities[5]. Residents are also being ordered to adopt personal precautionary practices including facemask wear in public areas by law.[6]

Earlier studies on the effectiveness of spread control measures during infectious disease pandemic showed large-scale strategies, such as closure of school closure, case isolation, household quarantine, internal travel restrictions and border control, were able to delay the spread and/or reduce incidence rate at certain periods through the outbreak season.[7–9]

Whilst awaiting the effectiveness of a series of measures to be seen, such evolving outbreak is expected to impose substantial burdens on healthcare system. Normally, a regional university-based hospital in China is equipped with 500-1,000 beds with only a small portion allocated for isolation purpose. Arising from the forecasting demands, increasing numbers of isolation beds and intensive care units (ICUs) for subsequent severe cases will be unquestionably required. Uncertainty of the capacity of current healthcare resources to tackle such sizable increase in demand is raised.

In anticipation of substantial burdens on healthcare system following this human-to-human transmissible epidemic, we aim to scrutinise the currently available information and evaluate the burden of healthcare system during the 2019-nCoV outbreak in China. We hope, by doing so, that the findings would be able to provide efficacious suggestions on reducing the spreads on the large scale and help authorities formulate effective control measures on combating this emerging viral outbreak.

## 2. Methods

In the classic SIR model, *S* represents the susceptible population, *I* represents the infected population, and *R* represents the recovered population. Susceptible population can be infected, who would be cured or died of the infection. The composition of susceptible, infected, recovered, deceased population is modelled based on a set of transition probabilities.

In this study, we applied a modified SIR model to evaluate the burden of healthcare system during the 2019-nCoV outbreak in Wuhan, China. Figure 1 shows the design of our model. Each cycle is one day in our model. The parameters used in the model were estimated based on the reported incidence released by the NHC of the People’s Republic of China or recent investigation on the outbreak (Table **1**). Daily reported incidence of confirmed 2019-nCoV cases, death, and recovery in China is available from 11^th^ January, 2020.[10] However, given that the probability of misdiagnosis is likely to be high in the early stage of the outbreak, we used the reported incidence between 0:00-24:00 on 28^th^ Jan, 2020 (the most updated data when the analysis was performed).

**Figure 1.**
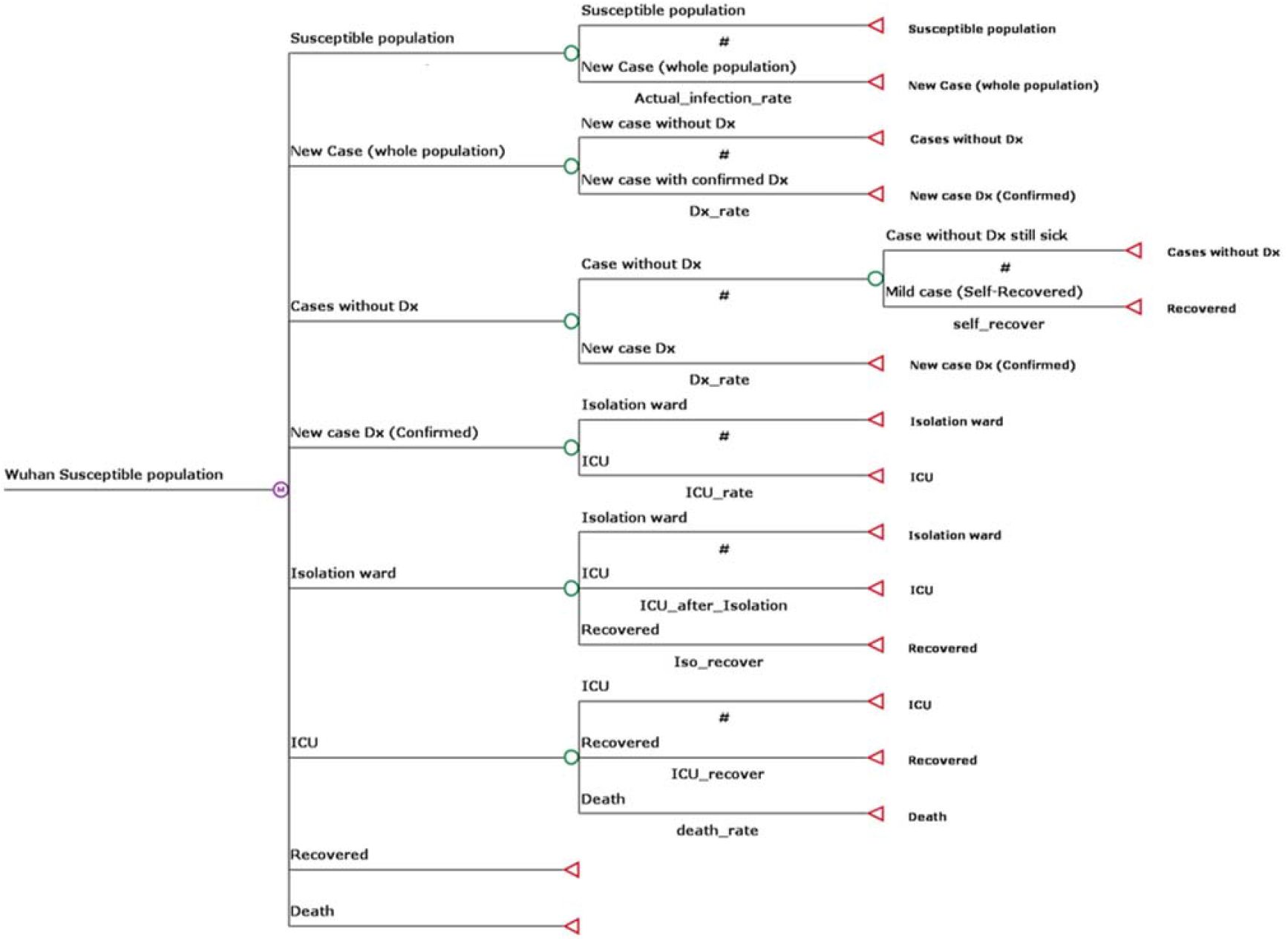
Design of the modified SIR model to evaluate the burden of healthcare system during the 2019-nCoV outbreak in China

**Table 1.** Parameter for the SIR model.

| Parameter | Interpretation<sup>a</sup> | Estimation<sup>b</sup> | Model input |
| --- | --- | --- | --- |
| Actual_infection_rate | Probability of being infected | Estimated based on data from National Health Commission of the People's Republic of China.[10]<br>$\text{Actual\_infection\_rate} = \frac{\sum(n\text{NewCase}/n\text{Population})}{n}$ | 0.013% |
| Dx_rate | Probability of being diagnosed if being infected | Assuming 10%, 50%, 90%, and 100% of the infected population can be accurately diagnosed. | 10%, 50%, 90%, 100% |
| self_recover | Probability of recovering if not being hospitalised | Assuming that cases without being diagnosed would recover in averagely 12.5 days. Thus, on any one day,<br>$\text{self\_recover} = \frac{1}{\text{Number of days needed to recover}}$ | 8.0% |
| ICU_rate | Probability of being admitted to ICU if being a confirmed case | Estimated based on data from National Health Commission of the People's Republic of China.[10]<br>$\text{ICU\_rate} = \frac{\sum(n\text{NewICU}/n\text{NewCase})}{n}$ | 18% |
| ICU_after_Isolation | Probability of being admitted to ICU after admitted to isolation ward | Assuming that the average number of days in isolation ward is 20 days and 5% of all cases admitted to the isolation ward would be transferred to ICU. Thus, on any one day,<br>$\text{ICU\_after\_Isolation} = \frac{5\%}{\text{Average number of days in isolation ward}}$ | 0.25% |
| Iso_recover | Probability of recovering if being admitted to isolation ward | Assuming that the average number of days in isolation ward is 20 days. Thus, on any one day,<br>$\text{Iso\_recover} = \frac{1}{\text{Average number of days in isolation ward}}$ | 5% |
| ICU_recover | Probability of recovering if being admitted to ICU | Assuming that the average number of days in ICU is 25 days. Thus, on any one day,<br>$\text{ICU\_recover} = \frac{1}{\text{Average number of days in ICU}}$ | 4% |
| death_rate | Probability of death if being infected and hospitalised | Assuming an overall death rate of 14% according to the recent investigation by researchers from the University of Hong Kong<sup>[11]</sup> and that the average number of days in ICU is 25 days. Thus, on any one day,<br>$\text{death\_rate} = \frac{14\%}{\text{Average number of days in ICU}}$ | 0.56% |

Abbreviation: intensive care unit (ICU)
<sup>a</sup> All probabilities are probabilities within one cycle (i.e., one day) in the SIR model;
<sup>b</sup> n is the number of days of which the data were used for estimating the parameter (n=1 in our analysis); nNewCase is the number of newly reported cases each day; nNewICU is the number of newly reported cases admitted to ICU each day; nPopulation is the number of population, in this analysis we used the number of population in Wuhan city, i.e., ~11 million; nNewRecover is the number of newly reported recovered patients each day;

The parameters included in the model were transition probabilities from one state to another within one cycle in the SIR model, i.e., one day. Briefly, we estimated the probability of being infected (Actual_infection_rate), and the probability of being admitted to ICU if being a confirmed case (ICU_rate) according to the reported incidence and the total population in Wuhan. Considering that cases without being diagnosed would mostly have mild symptoms, we assumed that the average number of days needed to recover to be 12.5 days for these cases. Thus, the probabilities of recovering if not being hospitalised within any given day (self_recover) was estimated by 1 divided by the average number of days needed to recover. We also assumed that, if being admitted to the hospital, the average number of days in isolation ward and ICU to be 20 and 25 days, respectively. Thus, the probabilities of recovering in any given day in the hospital were estimated by 1 divided by the average number of days in isolation ward (Iso_recover) or ICU (ICU_recover). We also assumed that a total of 5% of the confirmed cases admitted to isolation ward would experience deterioration of the symptoms and be transferred to ICU. Therefore, in any one day, the probability of being transferred to ICU from isolation ward (ICU_after_Isolation) was estimated by 5% divided by the average number of days in isolation ward. We considered an overall death rate of 14% among the hospitalised cases according to the recent investigation by researchers from the University of Hong Kong.[11] Therefore, the probability of being dead within any given day in the ICU (death_rate) was estimated by 14% divided by the average number of days in ICU. Lastly, the report by the MRC Centre for Global Infectious Disease Analysis at Imperial College London suggests there were a total of 4,000 cases of 2019-nCoV in Wuhan City (uncertainty range: 1,000 – 9,700) by 18th January 2020.[12] Comparing to the number of cases released by the NHC of the People’s Republic of China, this report suggests a diagnosis rate of less than 10%. Therefore, we considered multiple scenarios with different probabilities of being diagnosed (10%, 50%, 90%, and 100%) if being infected (Dx_rate).

## 3. Results

Figure 2 and Figure 3 depict our estimated daily numbers of beds occupied in isolation ward and ICU using a modified SIR model based on the currently available information. We generated curves of four hypothesised diagnosis rates to project the burden on healthcare system. For diagnosis rates of 100%, 90% and 50%, our projections showed that between 18,311 and 34,786 beds in isolation ward and between 4,938 and 9,346 beds in ICU would be needed by the end of January. However, in the scenario of 10% diagnosis rate, the predicted number of beds occupied is expected to soar and would reach 103,131 in isolation ward and 27,277 in ICU by the end of January.

**Figure 2.**
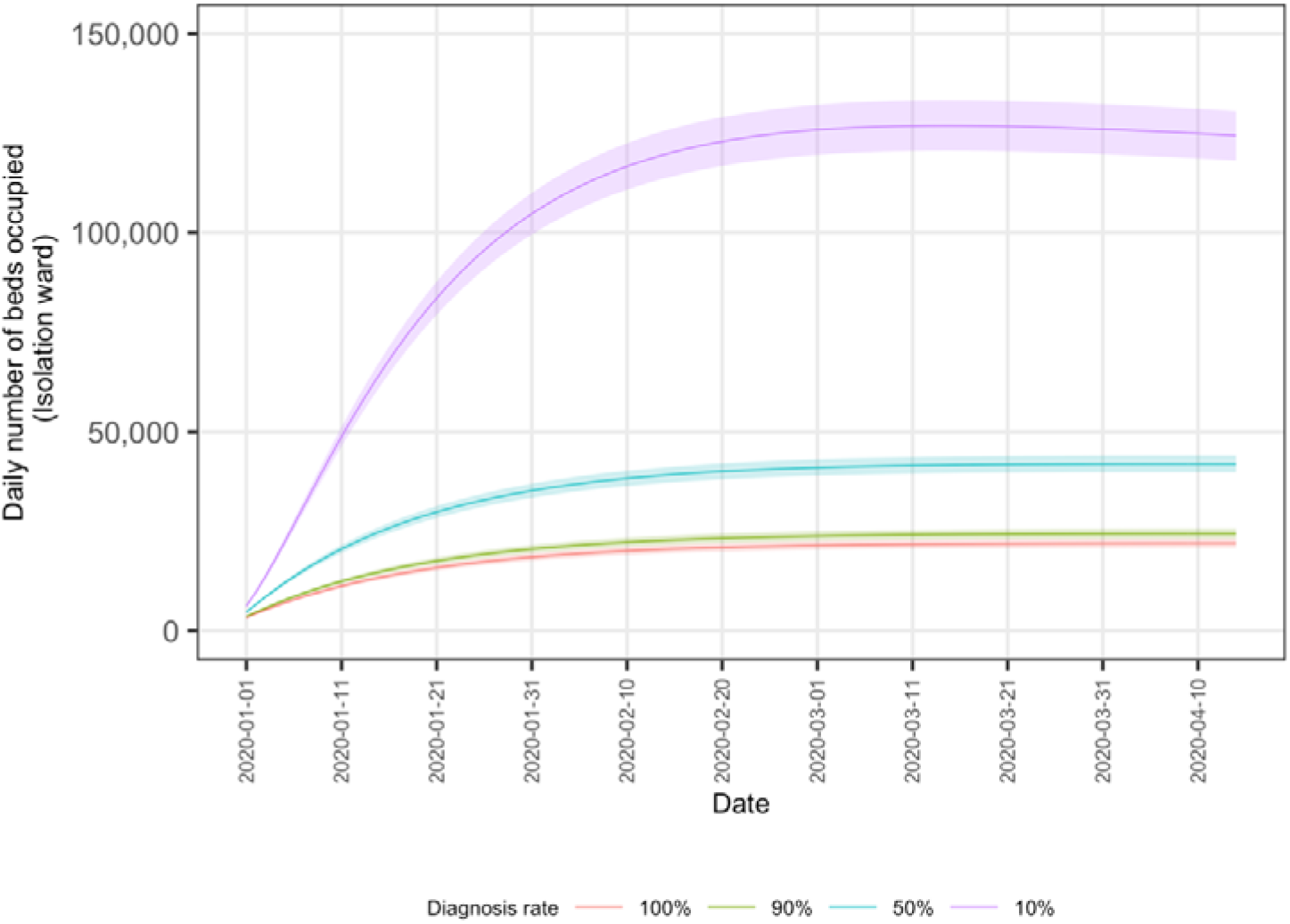
Estimated daily numbers of beds occupied (isolation ward) under scenarios with different diagnosis rates

**Figure 3.**
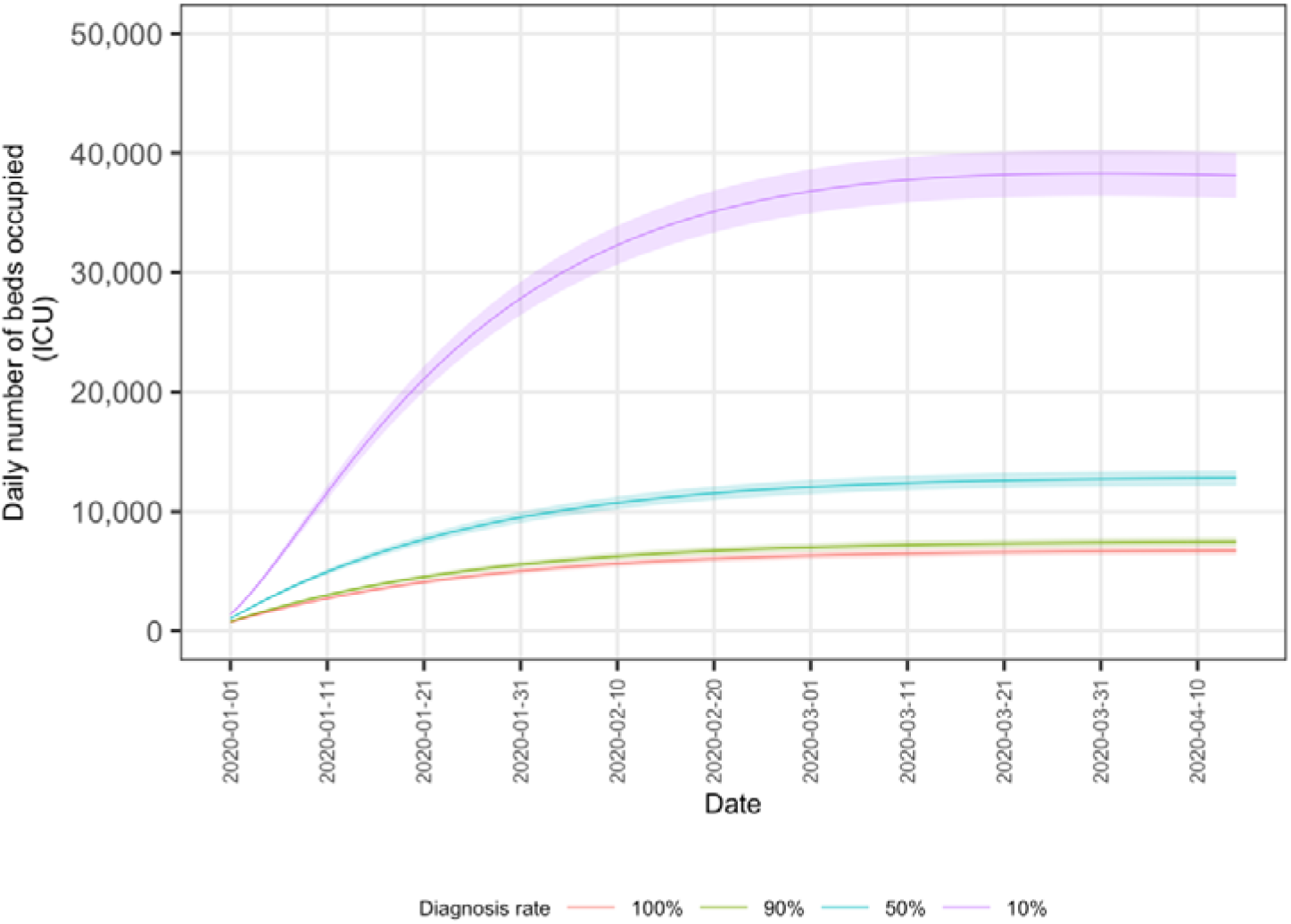
Estimated daily numbers of beds occupied (ICU) under scenarios with different diagnosis rates

We also investigated the specific burdens on isolation ward (Figure 4) and ICU (Figure 5) assuming a 50% diagnosis rate, given public health intervention scenarios of baseline (no intervention), 70%, 80% and 90% efficacy rates. If a 70% efficacy rate of public health intervention could be achieved, the number of cases being admitted to isolation ward and ICU would drop to a large extent throughout the course of outbreak. Similarly, greater benefits for healthcare system are expected to obtain if higher efficacy can be achieved (e.g., 80% or 90%). Total number of deaths would also be greatly reduced (Figure 6). By 31^st^ January, the total number of deaths under the no public health intervention scenario would be more than two times higher than that under the 70% efficacy rate of public health intervention.

**Figure 4.**
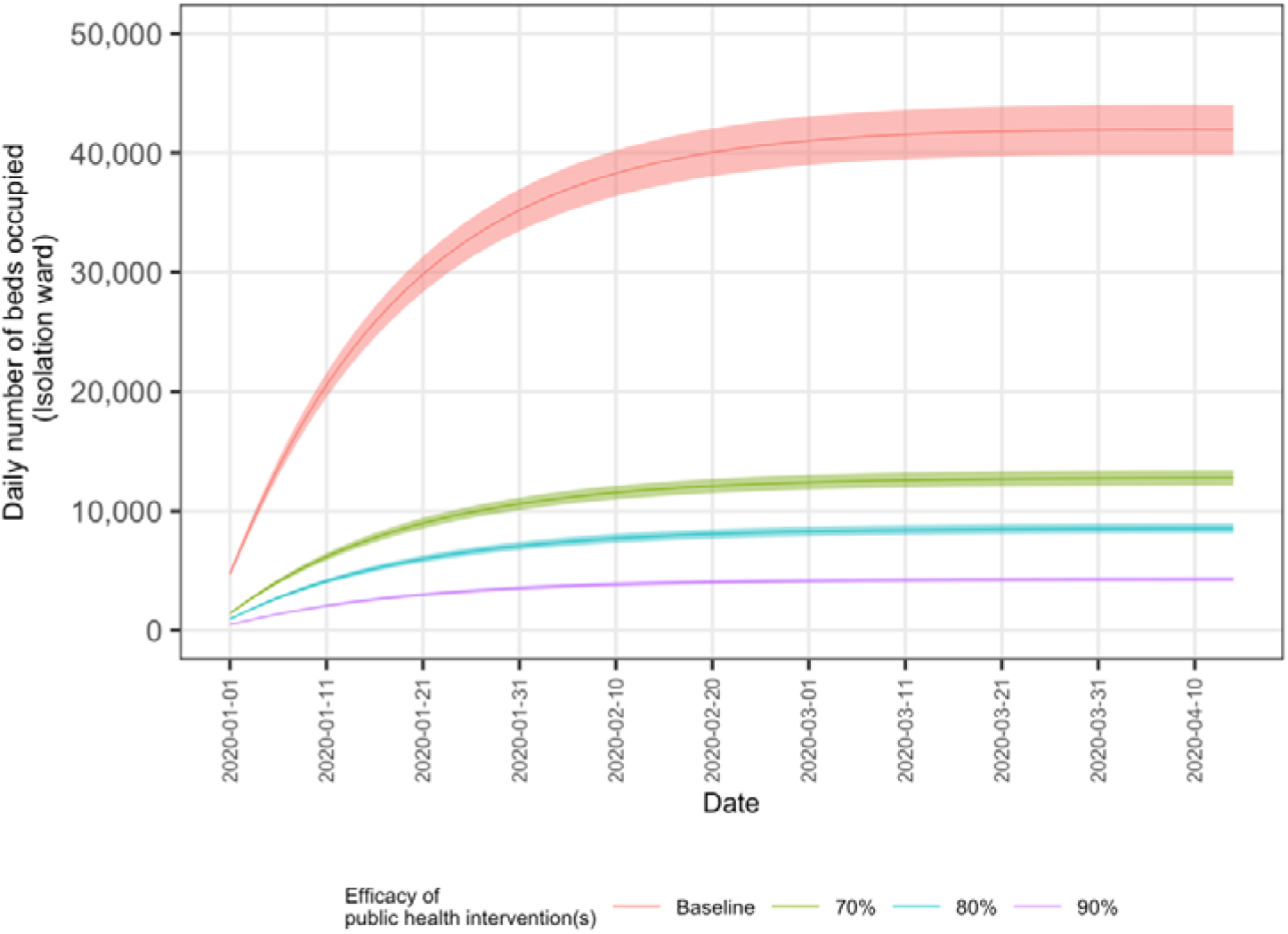
Estimated daily number of beds occupied (isolation ward) under different scenarios of public health intervention efficacy (assuming a 50% diagnosis rate)

**Figure 5.**
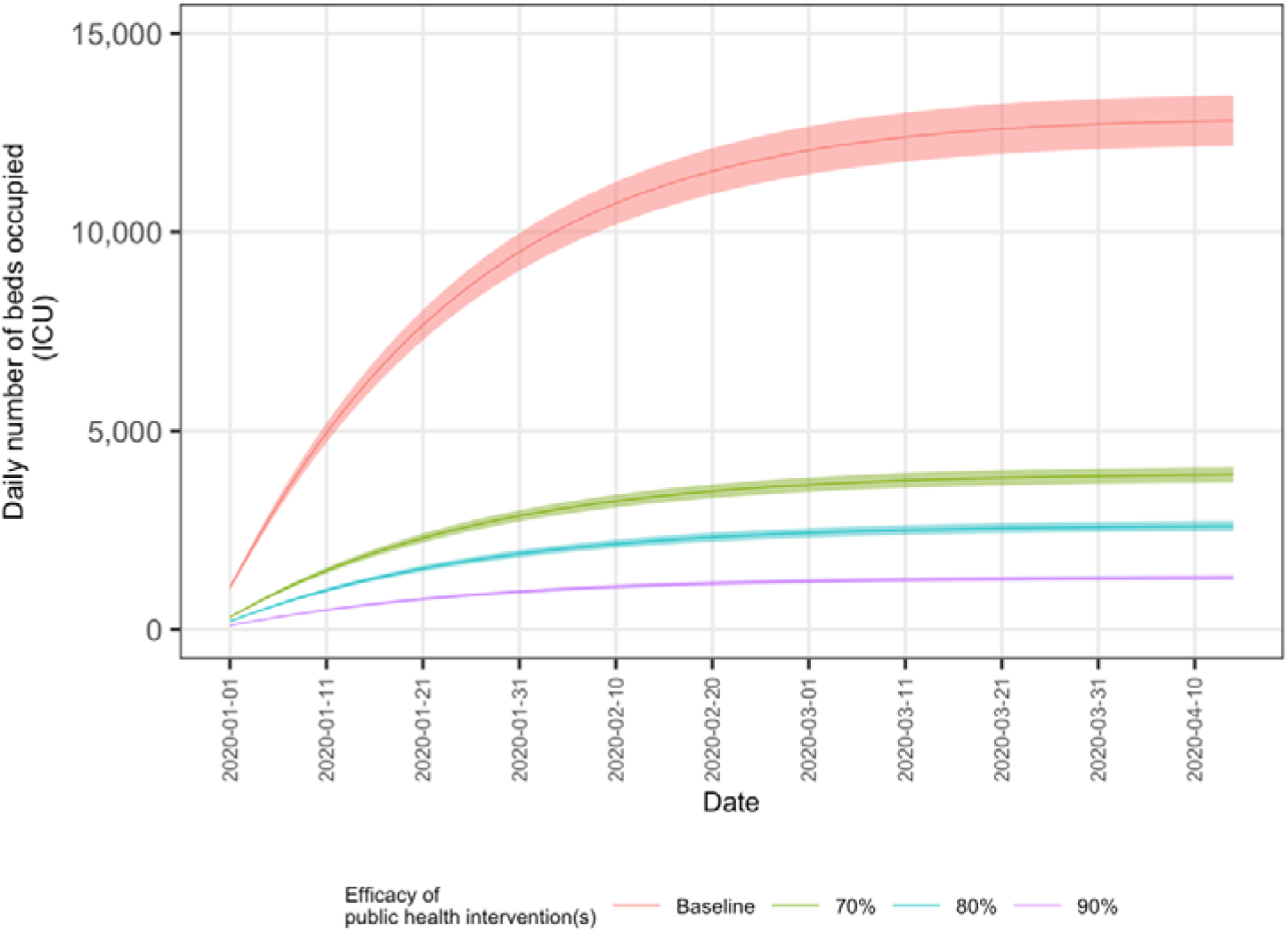
Estimated daily number of beds occupied (ICU) under different scenarios of public health intervention efficacy (assuming a 50% diagnosis rate)

**Figure 6.**
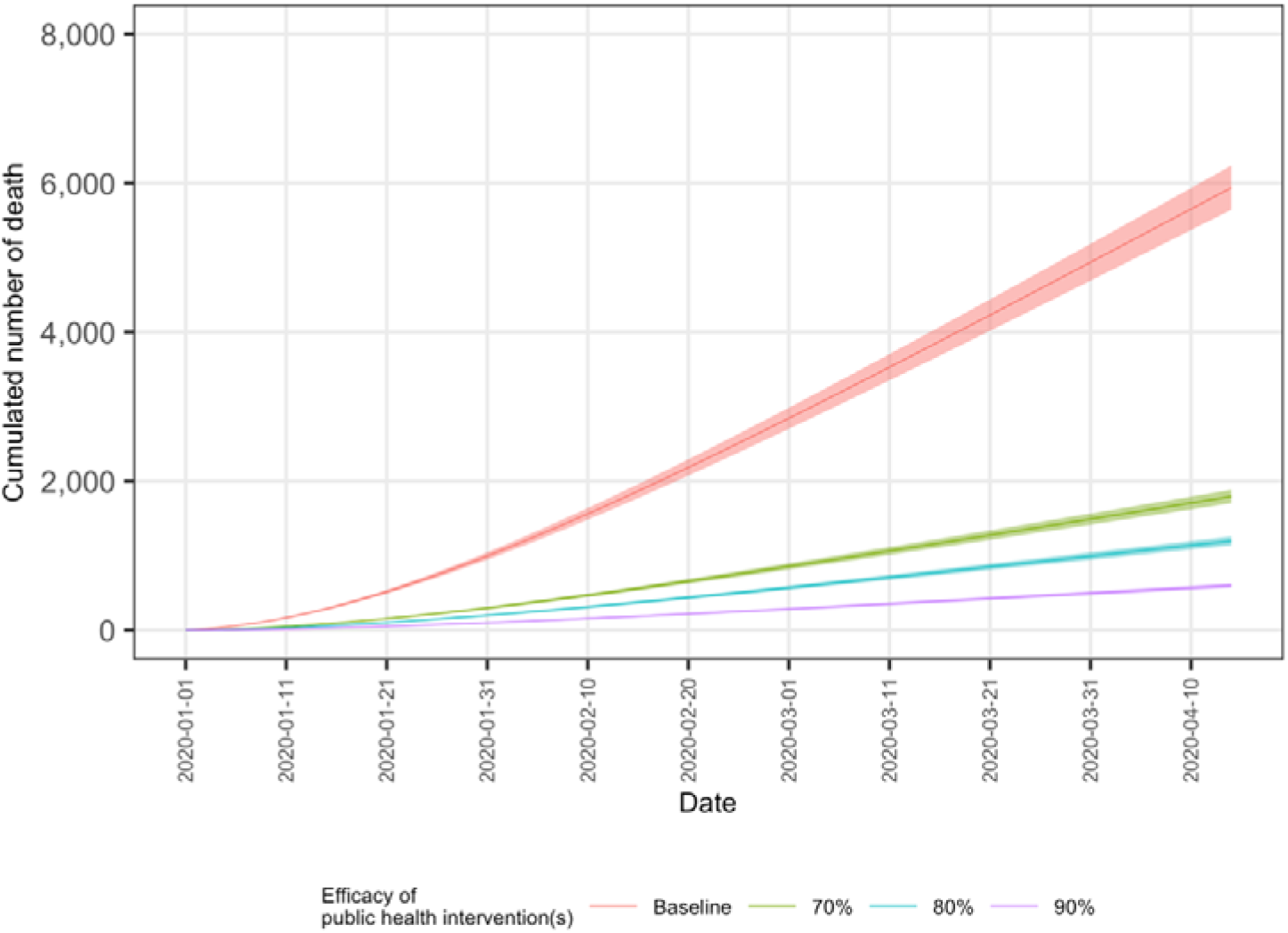
Estimated total number of deaths under different scenarios of public health intervention efficacy (assuming a 50% diagnosis rate)

As of 31^st^ January, it is estimated that there were 246,172 cases given a 10% diagnosis rate whilst being 88,075 and 52,094 cases given diagnosis rates of 50% and 90%, respectively, if no public health interventions were implemented (Table *2*). We further estimated the total number of cases with public health intervention efficacy of 70%, 80% and 90%, assuming a 50% diagnosis rate (Scenarios 4, 5 and 6). If 70% efficacy rate could be achieved (Scenario 4), the forecasting number of cases would drop dramatically to 11,056 as of 10^th^ February compared to 115,355 without public health interventions (Scenario 2). Even fewer cases can be expected if higher efficacy (e.g., 80% or 90%) is achieved.

**Table 2.** Estimated total number of cases in six different scenarios during the 2019-nCoV outbreak in Wuhan, China

|  | Scenario 1 | Scenario 2 | Scenario 3 | Scenario 4 | Scenario 5 | Scenario 6 |
| --- | --- | --- | --- | --- | --- | --- |
| Diagnosis rate | 10% | 50% | 90% | 50% | 50% | 50% |
| Public health intervention efficacy | No public health intervention | No public health intervention | No public health intervention | 70% | 80% | 90% |
| Estimated total number of cases (with undiagnosed mild cases) on 31<sup>st</sup> Jan 2020 | 246,172 | 88,075 | 52,094 | 26,498 | 17,672 | 8,840 |
| Estimated total number (with undiagnosed mild cases) of cases 10 days later (10<sup>th</sup> Feb 2020) | 331,524 | 115,355 | 67,883 | 34,736 | 23,170 | 11,591 |

## 4. Discussion

In this study, we synthesised available information during the 2019-nCoV outbreak occurring in Wuhan city, China, and estimated the burden on healthcare system effected by the increasing numbers of cases using a modified SIR model. Our results suggest the actual number of infected cases could be much higher than the reported as the spread continues. Therefore, the burdens on healthcare system would be substantial, particularly for the isolation wards and ICU, if no effective public health interventions were implemented.

Our analysis is limited by the availability of data and the lack of understanding of 2019-nCoV. First, SIR model can be substantially affected by the input parameters, i.e., the transition probabilities, which were estimated based on either the data released by the NHC of the People’s Republic of China or other recent investigations. Only the reported incidence between 0:00 and 24:00 on 28^th^ Jan 2020 was used at this stage. As more data become available, further investigation should be considered. Second, classic SIR model assumes a constant infection rate, which is not likely to be true as interventions being implemented. Therefore, in this study, we constructed SIR models with multiple efficacy rates of public health interventions as proxy for the change of infection rate. Third, our prediction can be influenced by the diagnosis rate. Therefore, we simulated a number of scenarios with different hypothesised diagnosis rates of 10%, 50% and 90% to estimate the actual number of infected cases. For the reason of mild or moderate symptoms reported by part of infected cases and the relatively long incubation period observed, it is reasonable to model a context of low diagnosis rate. In fact, recent investigation also suggests the current diagnosis rate could be low and likely lower than 10%.[13] Our estimates appear to be close to the recently published estimates[13].

Our results emphasised the vital importance of efficacious public health interventions during the course of outbreak. Established human-to-human transmissibility of this novel coronavirus can be one of epidemiological factors that contribute to the accelerated spreads within the epicentre Wuhan and towards cities and regions via transports of those with no or only subclinical symptoms. Sustained human transmission (i.e., basic reproduction number R_0_ > 1) is supported by the confirmed human-to-human transmissibility.[13] Therefore, upstream measures that limit or block the viral transmission between individuals within and across cities are urgently needed. Fortunately, the lockdown of Wuhan city has taken place and is believed to have largely minimised the spreads from the epicentre to other areas. However, the mayor of Wuhan later announced approximately five million of residents had left the city ahead of the implementation of lockdown (due to scheduled travel during Chinese new year and panic about the lockdown)[14], which may compromise the anti-spread effect of such city lockdown measure as to spreads to other cities. To date, information on transmission modes and severity of this novel coronavirus is still limited [15]. Further preventive measures to diminish contact between persons and reduce social distance, such as school closure, public transport shutdown, common activities suspension, etc.[7, 8], should be implemented as to avoidance of healthcare system breakdown.

To achieve higher efficacy of the public health interventions, efforts from individuals should not be neglected. In light of no available specific vaccines and treatments for such novel coronavirus thus far, a range of precautionary behaviours between individuals at homes and in communities are essential and vital to obtain proper control of the spreads in public and likely preventing superspreading events. Personal prevention strategies for seasonal influenza and other viral infections are still applicable during the present outbreak, inclusive of restricting ill residents from common activities, excluding symptomatic persons from entering homes/facilities, limiting visit especially of wet markets, live poultry markets or farms. Maintaining personal hygiene (e.g., frequently performing hand hygiene, washing hands with soap and water) and cough etiquette (e.g., covering nose and mouth when coughing, correctly disposing wasted tissues after coughing) are also beneficial.

Amongst these preventive practices, facemask wear appears to be the most operationalised and thus effective since it is observable from other members of the public. A recent cluster randomised controlled trial[16], consisting of multiple region-varying medical settings, showed both N95 respirators and medical masks can effectively prevent influenza and other viral respiratory infections. Apart from facemask wear being required in public of the epicentre city in Hubei province, it has also been mandated by the Guangdong provincial government that facemasks should be worn in public with effect from 26^th^ January, four days after the implementation in Wuhan. The Centre for Disease Control and Prevention, the United States, also emphasised the importance of wearing a facemask at all times when staying with infected individuals in shared spaces as one of precautions for large-scale spreads in community, specified in their interim guidance for prevention for 2019-nCoV from spreading in home and communities[17]. All these are extremely important in raising awareness in the public as to personal preventive steps given the present situation (mild or subclinical symptoms observed in many cases and observed long incubation period). Operational issues associated with wearing disposable facemasks to maximise their preventive effectiveness should be also publicised and educated to the general public including correct ways of wearing facemask, hygiene practices across the procedure of mask wearing, disposal of used masks.

Volunteering from healthcare professionals appears to play a major role in reduction of fears amongst the public. Imported healthcare support from other cities are certainly vital to the healthcare system in Wuhan during the present outbreak.[18] It is also noteworthy that volunteers with medical backgrounds (e.g., medical trainees, health science students) are actively instilling scientific insight and evidence-based knowledge via the Internet to eliminate rumours spread across Chinse social media [19]. We believe that these volunteering activities can contribute to a successful delivery of public health principle and, in turn, efficacious interventions.

The basic reproductive number of this novel coronavirus has been estimated in recent studies (range from 2.6 to 6.47)[13, 20–24], suggesting its spread being as similar as or more efficient than seasonal influenza. Our estimates also indicate that a country-wide outbreak or even an international outbreak is foreseeable. It is thus essential to implement effective public health measures to curb this very outbreak without delay. Otherwise, the current healthcare system would not be able to sustain. Once the breakdown occurred, mortality would be expected to soar due to lack of medical resources. Home isolation for patients with mild symptoms could be one of the possible managements if the ultimate limit of isolation beds at the city or province level was exceeded.

We hope that all the essential measures mentioned above can be inclusively implemented so as to achieve at least 70% efficacy as our projection. Any primary preventive steps are expected to be contributed to the curb on viral transmission and ultimately bring about the emergency situation to a controllable level via a functioning healthcare system.

To conclude, our estimates of the healthcare system burdens arising from the actual number of cases infected by the novel coronavirus appear to be considerable if no effective public health interventions were implemented. We call for continuation of implemented anti-transmission measures (e.g., lockdown of city, closure of schools and facilities, suspension of public transport) and further effective large-scale interventions spanning all subgroups of populations (e.g., universal facemask wear) with an aim to obtain overall efficacy with at least 70%-90% to ensure the functioning of and avoid the breakdown of healthcare system.

## Abbreviations

ICU: intensive care unit
NHC: National Health Commission

